# Csf1r-GCaMP5 Reporter Mice Reveal Immune Cell Communication in Vitro and in Vivo

**DOI:** 10.1101/2021.02.24.432710

**Authors:** N Taghdiri, D Calcagno, Z Fu, K Huang, RH Kohler, R Weissleder, TP Coleman, KR Kevin

## Abstract

Interconnected cells are responsible for emergent functions ranging from cognition in the brain to cyclic contraction in the heart. In electrically excitable cells, methods for studying cell communication are highly advanced, but in non-excitable cells, generalized methods for studying cell communication are less mature. Immune cells have generally been classified as non-excitable cells with diverse pathophysiologic roles that span every tissue in the body, yet little is known about their interconnectedness because assays are destructive and have low temporal resolution. In this work, we hypothesize that non-excitable immune cells are functionally interconnected in previously unrecognized cell communication networks. To test the hypothesis, we created a hematopoietic calcium reporter mouse (Csf1r-Cre × GCaMP5) and non-destructively quantified the spatiotemporal dynamics of intracellular calcium *in vitro* and *in vivo*. *In vitro*, bone marrow derived macrophages calcium reporters reveal that fatal immune stimulatory DNA-sensing induces rapid intercellular communication to neighboring cells. *In vivo*, using intravital microscopy through a dorsal window chamber in the context of MC38-H2B-mCherry tumors, Csf1r-GCaMP5 reporters exhibit spatiotemporal dynamics consistent with cell communication. We present a theoretical framework and analysis pipeline for identifying spatiotemporal locations of “excess synchrony” of calcium spiking as a means of inferring previously unrecognized cell communication events. Together, these methods provide a toolkit for investigating known and as-yet-undiscovered cell communication events *in vitro* and *in vivo*.

## INTRODUCTION

Interconnected cells underlie emergent functions, such as memory and cognition in the brain(1–3), metabolism in the liver(4), and cyclic contraction in the heart(5). Methods for investigating cell communication are well-established in electrically excitable cells such as neurons and cardiomyocytes; however, these represent less than 1% of all cells in the human body(6, 7). Considerably less is known about the connectedness of the > 99% of “non-excitable” cells because conventional assays are destructive or have low temporal resolution (8–10). The development of generalizable strategies for inferring cell communication in non-excitable cells would enable new cell communication networks to be discovered and characterized.

Cells communicate over diverse length and time scales, either by extracellular release and diffusive or convective transport, or by contact-dependent communication via mechanical coupling(11, 12), transmembrane signaling, or by direct exchange of cytosolic biomolecules via gap junctions (13). Such communication underlies normal physiological processes such as coordination of tissue morphogenesis during development, but it can also spread cell damage and innate immune activation in the context of tissue injury (14–18). Methods for studying cell communication in non-excitable cells in vitro include dye transfer following microinjection, scrape loading, gap-fluorescence recovery after photobleaching, or measuring electrical conductance via patch clamping (19). Calcium indicators have been loaded into cells and dynamically imaged to infer cell communication in cell culture (20–23) and in intact tissue slices (24). More recently, genetically-encoded calcium reporters have emerged to control and non-destructively measure membrane potential and calcium concentration, both in vitro and in vivo, without the need for loading exogenous dyes (25–30). These have been complemented by computational methods to infer communication in electrically excitable cells and to quantify calcium spikes in non-excitable cells; however, generalizable tools for inferring communication in non-excitable cells are in their infancy (31–37).

The immune system contains diverse non-excitable cells with only partially characterized roles within tissues. For example, at steady state, resident macrophages patrol local tissue microenvironments, combat pathogens, remove dead cells after injury, and even participate in unanticipated functions such as cardiac conduction(38, 39). In the context of cancer, immune cell communication underlies cancer recognition, antigen presentation, and cytotoxicity using a variety of fluorescent reporter systems (40). Yet, much remains to be learned about immune cell intercellular communication.

In this work, we test the hypothesis that immune cells are interconnected in previously unrecognized cell communication networks and that the dynamics of intracellular calcium can be used to recognize communication in non-excitable cells. Using the genetically encoded GCaMP5 reporter system driven by the hematopoietic selective Csf1r-Cre, combined with several computational pipelines, we discover unexpected immune cell calcium dynamics that enabled detection of cell communication both *in vitro,* after immunogenic double-stranded DNA stimulation, and *in vivo*, in the context of an orthotopic MC38 colon adenocarcinoma tumor. Together, the methods provide a generalizable toolkit for discovery and inference of cell communication in the Csf1r-GCaMP5 reporter specifically and across cell types, tissues, and disease states more generally.

## RESULTS

### Immune cell calcium reporter mouse construction and characterization

Resident immune cells influence diverse tissue processes across multi-cellular length scales, both in healthy tissue, and in diseases such as cancer. This led us to hypothesize that tissue immune cells are interconnected in unexpectedly dynamic cell communication networks. Since calcium is a highly dynamic second messenger affected by multiple signaling modules, it has high information content that can be used to recognize mutual information and information transfer in cell communities. Therefore, to enable non-destructive detection of spatiotemporally synchronous calcium dynamics in immune cells, we bred Csf1r-Cre mice with GCaMP5 calcium reporter mice and generated offspring in which the calcium reporter was expressed in all cells that expressed Cre recombinase under regulation of the Csf1r promoter (*SI Appendix*, Fig. S1*A-B*). The GCaMP reporter system consists of a genetically encoded calcium-dependent green fluorescence protein (GFP) as well as constitutive tdTomato reference protein(41). To characterize the distribution of reporter activation in the mouse, we performed confocal imaging of the heart, spleen, kidney, and lung (*SI Appendix*, Fig. S1*C*). We also performed flow cytometric analysis of peripheral blood leukocytes and of cells released from the digested solid tissues, which showed tdTomato+ cells in multiple hematopoietic lineages (*SI Appendix*, Fig. S1*D-F*). This allowed screening for dynamic calcium signals across a broad range of hematopoietic subsets and tissue resident immune cells.

### DNA Sensing precipitates Calcium dynamics of macrophage fatal immunogenic DNA sensing

We isolated bone marrow derived macrophages (BMDMs) from the Csf1rCre-GCaMP5 reporter mice, differentiated them in culture with m-CSF, and quantified their spontaneous activity across time in an environmentally-controlled live cell microscopy culture system. Time-lapse fluorescence imaging at a coarse sampling interval of 2 minutes revealed negligible spontaneous activation over 48 hours (*SI Movie* M*1*, Fig. 1*A*). Therefore, we introduced immunogenic dsDNA complexed with a transfection reagent, which is a spatially-localized innate immune stimulus capable of inducing cell communication and cell death via cGAS- and AIM2-dependent cytosolic DNA sensing pathways respectively (42–45). This revealed marked transient GFP signals as evidenced by maximum image projections, percent GFP+ area, and percent GFP+ cells (n=6, P<0.0001) (*SI Movie* M*2* Fig. 1*B-D*). We defined regions of interest and extracted single cell dynamics, which revealed asynchronous transient single cell GFP fluorescence spikes (Fig. 1*E-G*). The duration of each cellular calcium spike was self-similar (7 ±3 minutes per spike), but the spike initiation times were distributed across the 48-hour observation period at an initial rate of 24 spikes per hour, which declined to 9 spikes per hour by the final 2 hours. We observed no consistent spatiotemporal relationships between the location of cells and the timing of their calcium spikes (Fig.1*H*). Cells that exhibited calcium spikes at this sampling frequency did so only once, and detailed examination of the associated DIC images suggested the DNA stimulation precipitated cell death (Fig.1*I*), consistent with its ability to activate AIM2-dependent pyroptosis resulting in loss of cell membrane integrity and intracellular calcium overload (46). By quantifying loss of differential DIC variance across time, we were able to track the spatiotemporal dynamics of cell viability and show that immunogenic DNA stimulation induces sparse asynchronous cell death in BMDMs with single cell resolution (Fig.1*J*). This allowed investigation of how death of individual cells influenced surviving neighbor cells.

**Figure 1.**
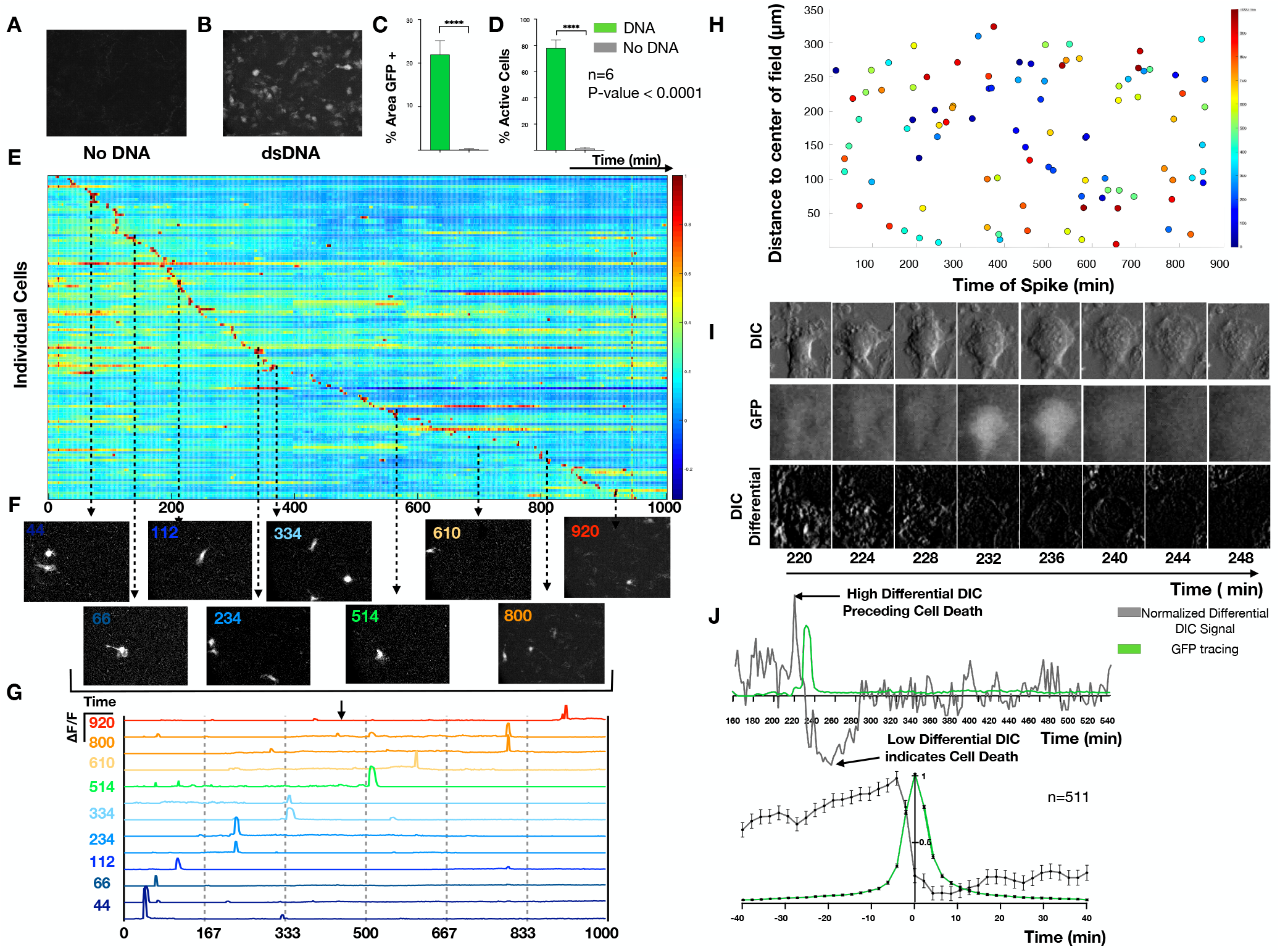
Calcium reporter macrophages reveal the dynamics of fatal DNA-induced calcium overload. Bone marrow derived macrophages (BMDM) were exposed to immune stimulatory dsDNA or vehicle control and live cell imaging was performed to quantify the resulting calcium signals every 2 minutes for 16 hours. A-B) Maximum intensity projections (MIPs) of representative time-lapse images for (A) no DNA compared to (B) dsDNA. C) Quantification of percent GFP+ area and D) percent active cells comparing no DNA to DNA (n=6). E) Heatmap of normalized calcium fluorescence versus time for each single cell over 16 hours. F) Example images at discrete time points illustrating asynchronous transient calcium reporter fluorescence. G) Quantification of the fluorescence dynamics of the cells shown in F. H) Spatial location of single cells with color indicating the timing of cellular calcium-dependent fluorescence spikes. I) DIC (top), GFP (middle), and differential DIC (bottom) surrounding a single calcium overload event. J) Example quantification of normalized differential DIC signal (grey) with low differential DIC indicating a lack of cell movement suggesting cell death. K) Stereotyped dynamics of differential DIC signals (grey) and calcium-dependent fluorescence (green) surrounding each calcium overload event (n=511).

### Macrophage cell communication is induced by fatal DNA-sensing

It is unknown to what extent cells sense the death of a neighboring cell. To investigate neighboring macrophage responses to DNA-induced cell death with high spatiotemporal resolution, we used scanning confocal microscopy at a sampling rate of 2Hz and confirmed that this did not undersample the GCaMP5 calcium dynamics (SI Appendix, Fig. S2) (SI *Movie* M3).

As above, steady-state BMDMs exhibited only rare spontaneous calcium spikes (*SI Movie* M4) while dsDNA stimulated BMDMs precipitated sparse asynchronous spatially isolated fatal calcium overload events evidenced by a sudden but sustained increase in high intensity GFP fluorescence that ultimately decreased as cell morphology transitioned to an inanimate “ghost” (*SI Movie* M5). Unexpectedly, this cell death precipitated dynamic fluorescence fluctuations in nearby bystander calcium reporter macrophages (*SI Movie* M*5*) (*SI Appendix*, Fig. S3*A-B*). We therefore tested the hypothesis that fatal-DNA sensing precipitates cell communication that can be inferred from temporal correlations of single cell calcium dynamics.

We developed a computational pipeline that converts raw time-lapse fluorescence images into single cell impulse trains representing the timing of calcium spikes. Briefly, single cell regions of interest (ROI) are defined, the background is corrected, single cell fluorescence versus time is extracted for each ROI, peak-finding is performed, and the resulting fluorescence peaks are transformed into impulses at the time of spike (TOS) for each peak. (Fig. 2*A-B*). Cells were sorted based on the timing of the maximum amplitude spike (TOS_max_) and plotted as a heatmap of single cell calcium dynamics, which revealed a temporal progression of spikes suggestive of intercellular signal propagation (Fig. 2*C*). Using the initial calcium-overloaded cell as a reference at time T0, we plotted the TOS_max_ for each cell versus its distance from the initiating cell, which revealed a propagation velocity of 9 μm/s (assuming a spherically symmetric diffusive model for propagation) (Fig. 2*D*), consistent with extracellular diffusion of a < 30kDa protein or restricted diffusion or intracellular transport of a lower molecular weight mediator. Inspection of individual time frames confirmed the isolated calcium overloaded cell precipitates a wave of non-fatal calcium spikes in neighboring cells (Fig. *E*). Cross-correlation of ranked spikes from the entire time-lapse recording predicted communication in the same temporal region (Fig. 2*F*). Taken together, these data reveal that cytosolic DNA sensing leads to sparse asynchronous isolated cell death that is rapidly communicated to neighboring cells where it precipitates nonfatal calcium dynamics.

**Figure 2.**
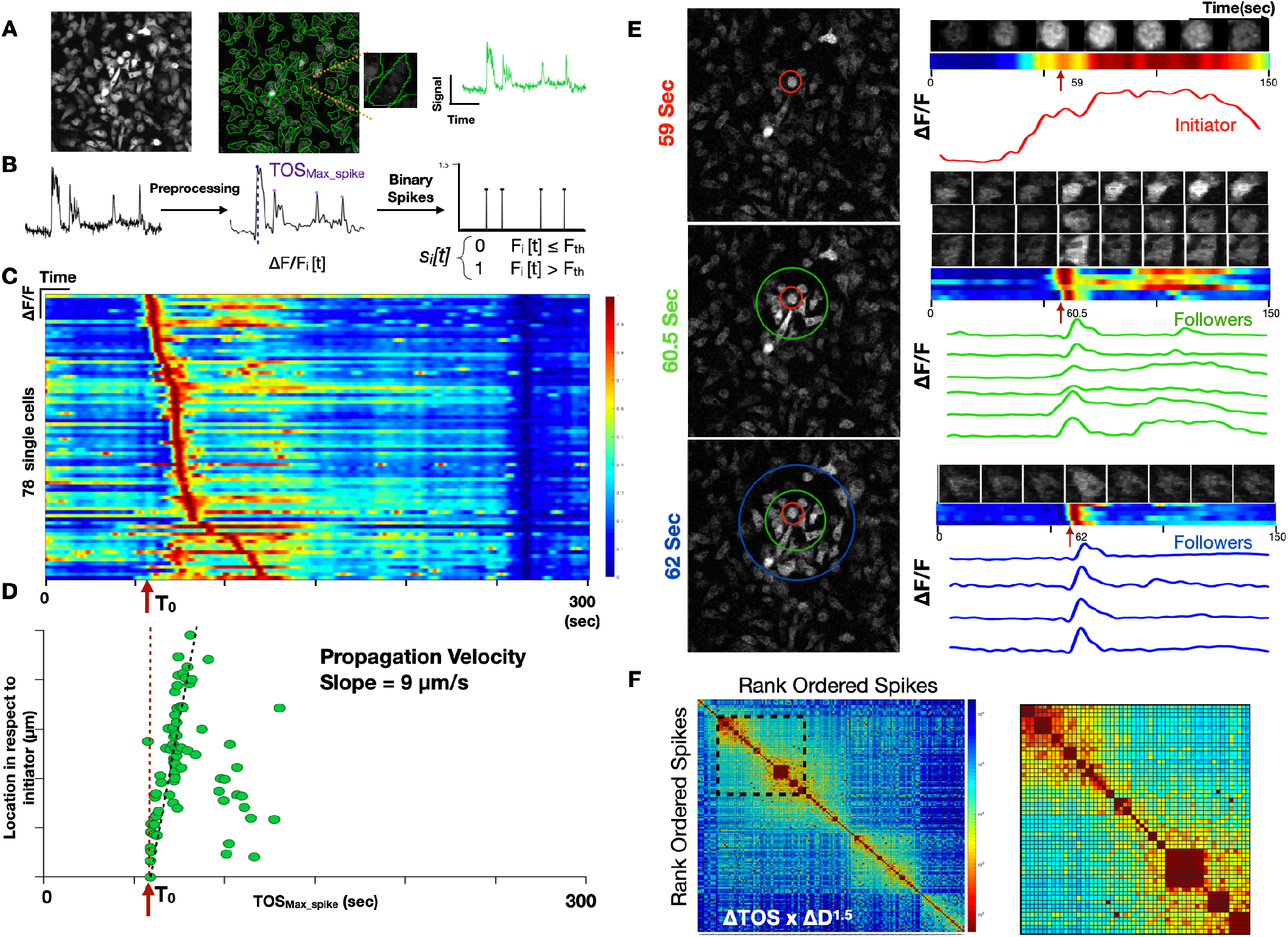
Macrophage DNA sensing induces cell communication in vitro. A) Maximum image projection of bone marrow derived macrophages derived from the Csf1r-GCaMP5 calcium reporter mouse (*left*). Regions of interest (ROIs) are drawn to define cell boundaries (*middle*). Individual cell with corresponding single cell fluorescence versus time tracing sampled at 2Hz (*right*). B) Signal processing including: Background correction, calcium intensity change quantification, bandpass filtering, peak-finding, determination of maximum spike(TOS_Max_spike_) based on the magnitude of calcium intensity and converting the fluorescence peaks into impulses at the time of spike (TOS) Signal processing to define spike times and convert to a binary impulse train. C) Heatmap of single cells (y axis) versus time where intensity represents normalized change in fluorescence ΔF/F. The time of the initiating cell spike is defined as T0. D) Distance of cell from the initiating cell versus time of maximum spike intensity. This reveals a slope of 9μm/s which is interpreted as the communication propagation velocity. E) Frames of the time lapse fluorescence during the propagation are shown at 59s, 60.5s, and 62s (left). Color coded concentric rings are shown to define the initiator cell, exhibiting sustained fluorescence (red) and the secondary responders (green and blue concentric rings), exhibiting brief calcium spikes. At right, time-lapse montage of individual cell calcium spikes are shown with the corresponding heatmap and fluorescence versus time tracing. F) Cross-correlation heatmap of rank ordered spikes for all cells. Color represents the product of spike time difference and Euclidean spatial distance ΔTOS × ΔD^1.5^. Inset shows the region of high cross-correlation indicating spikes that are highly localized in time and space.

### Inference of cell communication from Csf1r-GCaMP5 reporter dynamics *in vivo*

Next, we asked whether the calcium reporter would enable inference of cell communication *in vivo*. We installed a dorsal window chamber in the Csf1r-GCaMP5 reporter mouse and performed time-lapse imaging (Fig. 3*A*, *SI Movie* M*6*). We implanted 1 million MC38-H2B-mCherry colon adenocarcinoma cells into the tissue underlying the window chamber and observed a dynamic host response. Within 24-72 hours of MC38-H2B-mCherry cell implantation, we discovered host Csf1r-GCaMP5 reporter cells with highly dynamic calcium spikes. We defined ROIs and quantified the single cell fluorescence dynamics of each cell across time. Heatmaps of the normalized fluorescence from hierarchical unsupervised clustering suggested that some cells exhibit highly synchronous calcium spikes amidst a background of seemingly random asynchronous calcium spikes (Fig. 3*B*). Inspection of cross-correlations of rank-ordered calcium spikes revealed the same timing and duration of calcium spike synchrony (Fig. 3*C*). We therefore set out to determine if the observed synchrony was best explained by cell communication.

**Figure 3.**
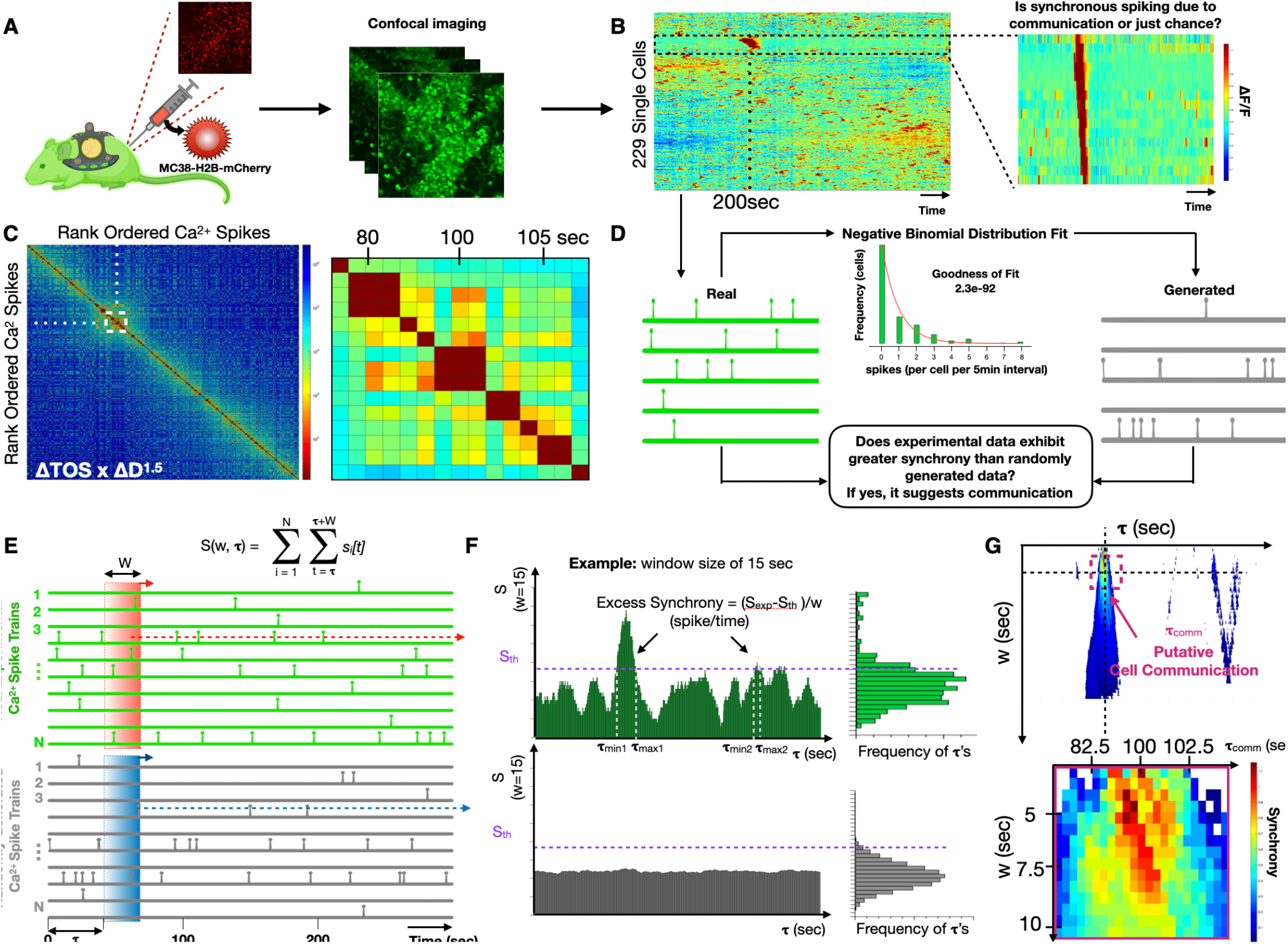
Csf1r-GCaMP5 reporter cell communication in vivo revealed by intravital imaging through a dorsal window chamber. A) Cartoon illustration of the intravital imaging of a calcium reporter mouse with a dorsal window chamber and orthotopically injected MC38-H2B-mCherry tumor cells. B) Heatmap of single cell fluorescence dynamics hierarchically clustered with single cells (y axis) versus time (x axis) where color represents the normalized change in fluorescence ΔF/F. Inset shows a cluster of cells with temporally localized calcium spikes. C) Cross-correlation heatmap of rank-ordered calcium spikes for all cells. Color represents the product of spike time difference and Euclidean spatial distance ΔTOS × ΔD^1.5^. Inset shows the region of high cross-correlation indicating spikes that are highly localized in time and space. D) Strategy for constructing “generated” (simulated) cells sampled from the same negative binomial distribution of spike trains as the “real” experimental spike trains. Comparisons allow estimation of whether synchrony occurs by chance or exhibits “excess synchrony”, beyond chance, which we interpret as putative cell communication. E) Method for quantifying normalized number of spikes (*S/w*), also known as “synchrony”, for *real* and *generated* single cell spike trains, where *S* is a function of temporal window size, *w*, and window initiation time, *τ*. F) Method for defining the timing of “excess synchrony” of *real* cell populations compared to the corresponding *generated* populations. G) Heatmap of excess synchrony (ΔS/w) as a function of temporal window size, *w*, and window initiation time, *τ*. Inset shows timing of high excess synchrony and putative cell communication.

### “Excess Synchrony” – a metric for identifying cell communication events in vivo

To determine if correlated calcium spiking is due to cell communication or if it can be explained simply by chance, we defined a metric called “excess synchrony”, which quantifies the extent to which calcium spikes in a specific window of time exceed what would be expected from randomly generated spikes sampled from a population of statistically-comparable synthetic cells. We first used the experimental data to assemble the overall distribution of spike frequencies (spikes per 5-minute recording) and modeled it as a negative binomial (goodness of fit of 2.3×10^−92^). Next we sampled from the distribution to create a “generated” population of synthetic calcium spiking cells (Fig. 3*D*). Synchrony was then defined as the number of spikes (*S*) within a defined region of time (***τ***), and *excess synchrony* was defined as (*ΔS/w),* the difference between experimental *S_exp_* and S threshold (S_th_), the 95^th^ percentile of the generated data, normalized to the temporal window size *w* (Fig. 3*E*). At the extremes of large and small window sizes, non-communicating spikes dominate and *ΔS* approaches zero; however, at an optimal window size (*w_opt_*), a putative cell communication process creates a local maximum of *excess synchrony (ΔS/w)* due to temporally concentrated calcium spikes that exceed the synchrony predicted by generated cells (Fig. 3*F*). By plotting a heatmap of excess synchrony ΔS/w = (S_exp_ - S_th_)/*w* as a function of window size and region of time, we were able to identify the duration of timing of communication (***τ _comm_***) by determining the 80th percentile of maximal excess synchrony’s timing from each window size. Accordingly, in order to find the putative cell communication, the optimal window size (*w_opt_*) was recognized by investigating the maximal excess synchrony window size within the duration of timing of communication. (Fig. 3*G*).

To validate the predictions, we examined the spatial distribution of synchronous cells at time **τ** and observed two qualitative groups, one that was highly localized in space and one that was disperse (Fig. 4*A*). We separated these populations using unsupervised k-means clustering of Euclidean distances (Fig. 4*B*). Heatmaps of normalized fluorescence dynamics showed that localized cells from cluster 1 only spiked 1-2 times during the recording whereas spatially disperse cells from cluster 2 spiked more frequently and likely represented the cells that were synchronous “by chance” as predicted above (Fig. 4*C-D*). Inspection of the time-lapse images confirmed these predictions, as it showed organized wave of calcium fluorescence propagation in cluster 1 but no qualitative evidence of communication in cluster 2 (Fig 4*E-G*). Cluster 1 also exhibited significantly higher calcium fluorescence full-width half-max (FWHM) and relative calcium spike amplitude (ΔF/F) (Fig. 4*H-K)*. Together, these data demonstrate that the calculation of “excess synchrony” of calcium spikes can be used to identify cell communication events *in vivo* amidst a background of random calcium spikes.

**Figure 4.**
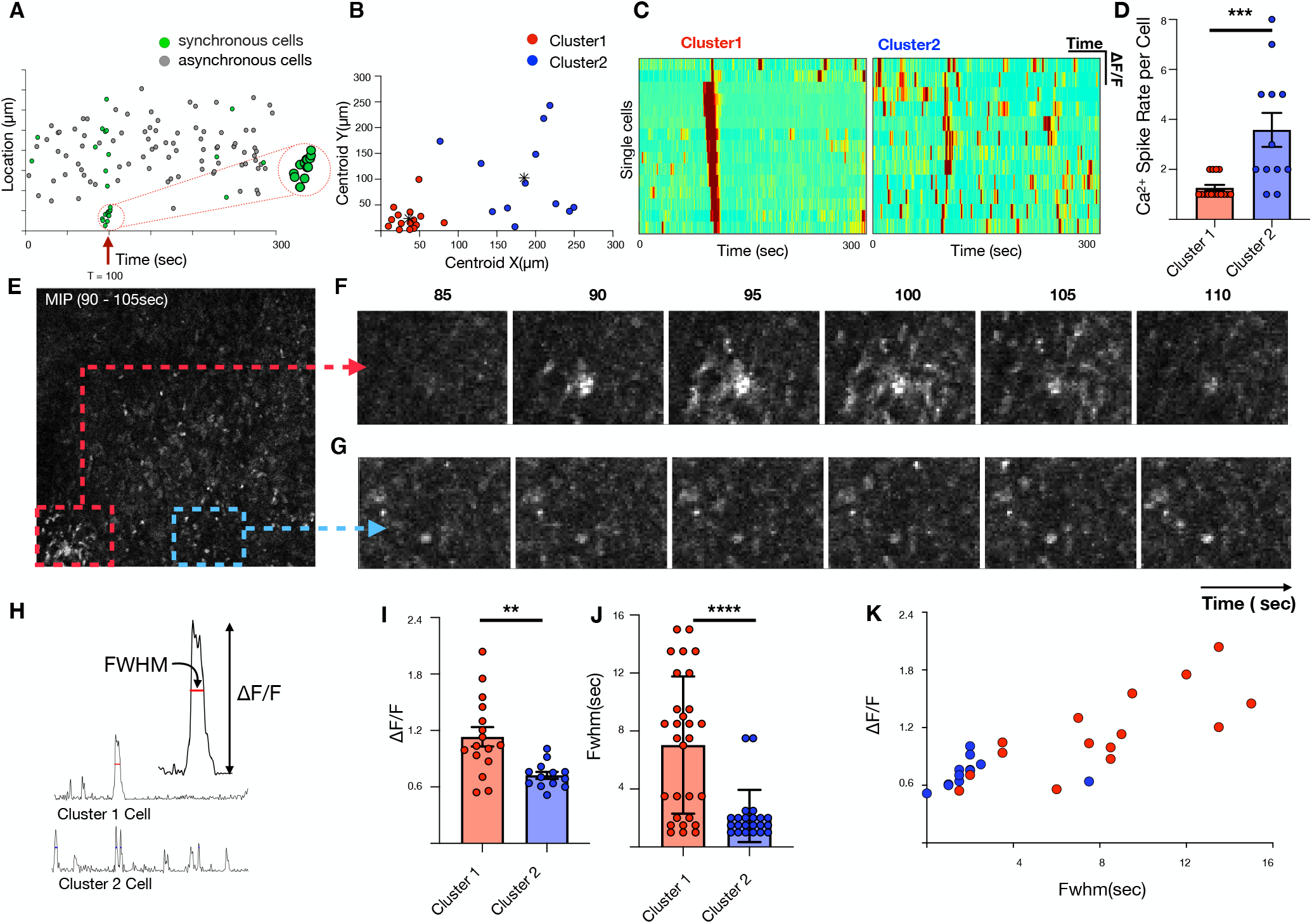
Characterization of spatiotemporally synchronous spiking cells. A) Correlation between location and timing of individual cells that spike during synchrony event. B) Unsupervised k-mean clustering based on Euclidean distances reveals 2 categorical clusters of synchronous cells – those that are spatially localized (red) and those that are disperse (blue). C) Heatmap of single cell fluorescence dynamics for Clusters 1 and 2. D) Comparison of single cell calcium spike rate for Clusters 1 and 2. E) Maximum intensity projection spanning the time of high excess synchronicity (90-105 sec). F-G) Localized region containing Cluster 1 cells (F) compared to comparable-sized region (G). H-K) Comparison of normalized calcium-dependent fluorescence changes (ΔF/F) and full width half max (FWHM) of Clusters 1 and 2.

### Prediction of spatiotemporal cell communication and regulation in a tumor context *in vivo*

To accommodate regions of high calcium spike frequency, we expanded the temporal synchrony pipeline to add spatial resolution. A maximum image intensity plot of such a recording is shown (Fig. 5*A,* SI *Movie* M*7*). In this data set, the dynamics of 522 single cells were quantified using the pipeline above and displayed as a heatmap (Fig. 5*B*). We spatially subset the field into 50μm × 50μm sub-images with 25 μm overlap to identify local areas of excess synchrony (*SI Appendix*, Fig. S4*A-B*). This resulted in a 3D volume of excess synchrony versus space and time (Fig. 5*C*), which enabled identification, localization, and counting of cell communication events at different levels of stringency. Slices from the volume at locations of excess synchrony are shown as a function of space (Fig. 5*D, E*), as a function of time (Fig. 5*F, G*), and as a kymograph in space-time (Fig. 5*H,I*). We repeated the analysis for several temporal window sizes and synchrony stringencies to show how predicted cell communication events would vary (Fig. 5*J*). The resulting self-similar curves revealed peaks of synchrony when the window size w approximated the characteristic time scale of cell communication. Increasing the stringency of the synchrony threshold leads to fewer but more prominent cell communication events as expected. Finally, we compactly displayed the results by color-encoding synchrony events on a spatial map to facilitate validation compared to the underlying fluorescence time-series data (Fig. 5*K*).

**Figure 5.**
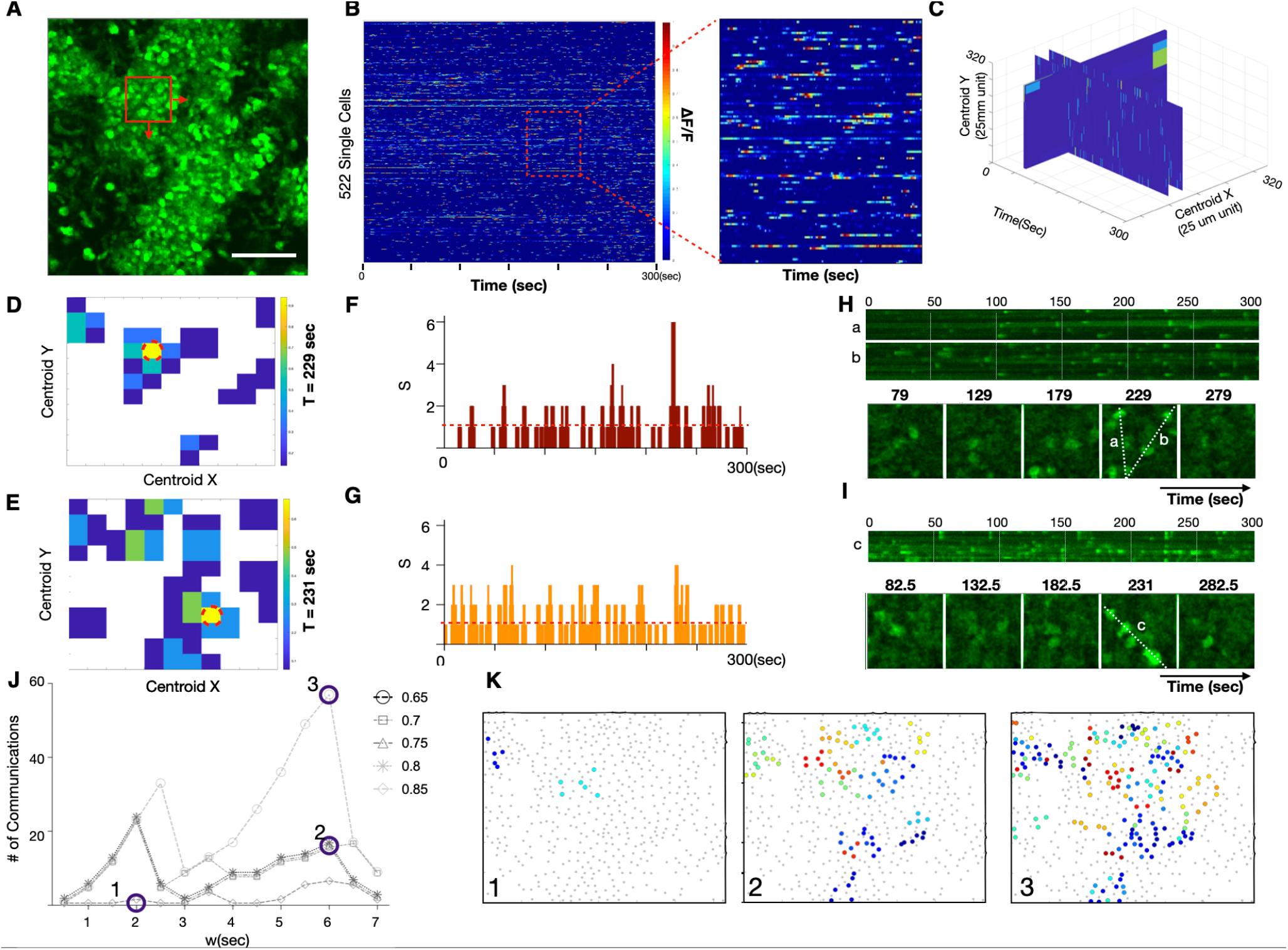
Quantification of excess synchrony and inference of communication in highly dynamic in vivo microenvironments. A) Maximum intensity projection of highly dynamic cellular population in vivo. (scale bar = 80μm) B) Heatmap of 522 single cell calcium fluorescence dynamics versus time. Color represents normalized calcium-dependent fluorescence changes ΔF/F. Inset shows close-up of individual cells. C) Spatiotemporal calcium dynamics as a volume of 50um × 50um regions shifted by 25um increments in the x and y directions over the 300 second (600 sample) time-lapse recording. D-E) Two time points where excess synchrony is calculated versus x and y location. The red dotted circle indicates the location of the highest excess synchrony. F-G) Number of spikes (S) versus time for the high synchrony locations circled red in D and E. The threshold for treating S as excess synchrony and considering it a putative cell communication event is shown as a red horizontal dotted line. This line can be increased to create a more stringent threshold for interpreting synchrony as putative cell communication. H-I) Line scans (top) and montage of individual time frames (bottom) for the locations circled red in D and E. J) Number inferred communication events as a function of temporal window size and synchrony stringency. K) Locations of communication events at different window sizes and synchrony stringency. Color indicates the timing of peak communication (maximum excess synchrony).

## Discussion

We have described construction and characterization of a Csf1r-GCaMP5 calcium reporter mouse that reveals unexpected dynamics and communication between immune cells *in vitro* and *in vivo. In vitro*, the reporter and the associated analysis pipeline enabled direct visualization of fatal calcium overload precipitated by DNA sensing and rapid non-fatal communication to surviving macrophage neighbors. *In vivo*, in the context of an MC38-H2B-mCherry tumor (SI *Movie* M*8-9)*, it enabled immune cell communication to be inferred from spatiotemporal analysis of Csf1r-GCaMP5 calcium spikes. Because cell communication occurs in a background of ambient calcium fluctuations of unknown etiology, we defined a metric termed “excess synchrony” and an associated computational pipeline to identify putative cell communication events amidst ambient random calcium spikes. The tools are highly generalizable and should be applicable to the Csf1r-GCaMP5 as well as other genetically encoded calcium reporter experiments, both at steady-state and in the context of disease.

Calcium is not unique to a single signaling pathway. Nevertheless, calcium reporters are among the most dynamic genetic reporters available, which allows them to non-destructively monitor cellular conversations and delineate which cells are “talking”, when, where, for how long, and at what volume and tempo. This is akin to watching a movie in a foreign language. Although one does not know the meaning of the words, one can readily determine which characters are conversing, when, where, to whom, in what context, and with what apparent content and emotion. With enough observations, one can gain insight into the relationships between the characters and the dynamics of their interactions despite not knowing the vocabulary, just as one can infer communication from correlated intracellular signaling without knowing the precise upstream biological inputs or signal transduction pathways.

This work provides an important addition to the immunologist’s toolkit, which currently consists of fluorescent reporters primarily designed to measure cell location, migration, or dwell time. The Csf1r-GCaMP5 calcium reporter not only enables measurement of dynamics, but it enables inference of cell communication aided by the computational pipeline and the calculation of “excess synchrony”. Future studies will explore what information these calcium signals encode, and upon what molecular mechanisms the associated intracellular communication depends. This can be approached by examining genetic or pharmacologic modulators of candidate communication molecules and measuring whether observed calcium spike synchrony is altered.

In this study, we used the new reporter to examine cell communication in two biological contexts. Cytosolic DNA sensing is a critical step in pathogen and cancer cell recognition. It can lead to pyroptotic cell death and can precipitate gap junction-mediated transfer of second messengers; however, such communication has previously been inferred from end-point assays of co-culture experiments or measurements of downstream signaling. Here, the Csf1r-GCaMP5 reporter enabled previously unobserved rapid DNA-induced intercellular communication to its surviving neighbors on the time scale of seconds. The fact that cells become rapidly aware of neighboring cell death is intriguing and motivates future experiments to explore what downstream signaling and phenotypic consequences this may trigger. Doing so will likely require genetic modification of candidate mediators and/or receptor machinery. Chimeric approaches that combine the reporter with genetically manipulated cells or mice are likely to be particularly illuminating when dissecting candidate communication mechanisms. In addition, spectrally diverse calcium indicators will enable non-destructive multi-color calcium dynamics to report on two-way communication between host and tumor, thus providing a vivid picture of the intercellular conversations within tumors and in other complex tissues.

In conclusion, we describe a new addition to the experimental immunologist’s toolkit. We show that calcium is a dynamic and high information-content mediator that can be non-destructively monitored in cell communities to infer previously unrecognized intercellular communication networks. As genetically encoded calcium indicators continue to advance and diversify, they will shed light on myriad undiscovered cellular conversations, not just limited to the brain, but across every organ, both during health and disease.

## Acknowledgements

The work was funded by NIH R00HL129168 (K.R.K.) and DP2AR075321 (K.R.K.), 5R01HL122208 (R.W.), 5R01HL131495 (RW), 5R01CA206890 (RW)

## Author Contributions

N.T. and K.R.K. designed the study. N.T., D.C, F.Z., K.H. R.K., and K.R.K. performed experiments. All authors analyzed data and synthesized results. N.T. and K.R.K. wrote the initial draft and all authors edited the manuscript.

## MATERIALS AND METHODS

### Construction of the immune cell calcium reporter mouse

Mouse experiments were approved and conducted under the oversight of University of California San Diego Institutional Animal Care and Use Committee (#17144) or approved by the Subcommittee on Animal Research Care at Massachusetts General Hospital. All mice were maintained in a pathogen-free environment. Csf1r-GCaMP5 calcium reporter mice were created by breeding “Csf1r-Cre” C57CL/6-Tg(Csf1r-cre)1Mnz/J) (The Jackson Laboratory; stock 029306) with “GCaMP5” calcium reporter mice, B6;129S6-Polr2a^Tn(pb-CAG-GCaMP5g,-tdTomato)Tvrd^ (The Jackson Laboratory; stock 024477) to generate offspring in which the calcium reporter was expressed in all cells that expressed Cre recombinase under regulation of the Csf1r promoter (41, 47). Genotyping was performed per Jackson Laboratory instructions.

### Tissue processing

Peripheral blood for flow cytometric analysis was collected by retro-orbital bleeding using heparinized capillary tubes (BD Diagnostic Systems) and red blood cells were lysed with 1x red blood cell lysis buffer (BioLegend). For organ harvest, mice were perfused through the LV with 10 mL of ice-cold PBS. Hearts, spleen, lung and kidney were enzymatically digested for 1h under continuous agitation at 37C in 450 U/ml collagenase I, 125 U/ml collagenase XI, 60 U/ml DNase I, and 60 U/mL hyaluronidase (Sigma) and filtered through a 40μm nylon mesh in FACS buffer. to generate a cell suspension for staining and flow cytometric analysis as previously described (48). To define the anatomical distribution of Csf1r-Cre-induced reporter cells within solid organs, we cut 1mm sections with a tissue slicer (Zivic Instruments) and examined the spatial distribution of reporter fluorescence in each tissue using a Nikon STORM super resolution confocal microscope at UCSD.

### Flow cytometry

Isolated cells were stained at 4°C in FACS buffer (PBS supplemented with 2.5% bovine serum albumin) with and hematopoietic lineage markers including Ly6G (BioLegend, clone 1A8, 1:600), CD11b (BioLegend, clone M1/70, 1:600) and Ter119 (BioLegend, clone TER-119, 1:600). This was followed by a second staining for NK1.1 (BioLegend, clone PK 136, 1:600), Thy1(CD90.2, BioLegend, clone 53-2.1 1:600), and Ly6C (BioLegend, clone HK1.4 1:600). Cell suspensions were labeled with DAPI just prior to flow cytometric analysis to allow exclusion of dead cells.

Doublets, erythrocytes, and dead cells were excluded by forward scatter, Dapi, and Ter119. Neutrophils were identified as (Ter119^low^/CD11b^high^/Ly6G^high^). Monocytes were identified as (Ly6G^low^/Ter119^low^/CD11b^high^/Ly6C^high^). NK or T cells were identified as (CD11b^low^/Ly6G^low^/(Nk1.1^high^ or CD90.2^high^)) respectively. The Cre-induced fraction of each hematopoietic lineage subset was determined based on the fraction that was tdTomato^high^. Data was acquired by Sony sorter MA900 at UCSD and analyzed with FlowJo software.

### Cell Culture

Bone marrow derived macrophages (BMDMs) from the Csf1r-GCaMP5 reporter mice were isolated, cultured in 10% FBS 1% Pen/Strep-containing DMEM, and differentiated with addition of 10ng/mL recombinant m-CSF (Peprotech) (every other day media changes) for a period of 7 days as previously described. 10μg of immunogenic HT DNA (Invivogen) was complexed with a Lipofectamine transfection agent (ThermoFisher) in serum free-media and added to 1 million BMDMs in a 6-well multi-well plate with serum- and mCSF-containing media for each experiment.

### Time-lapse Imaging of Calcium Reporter Dynamics in Vitro and in Vivo

Low temporal frequency imaging was performed using an Olympus Vivaview epifluorescence microscope with a sampling interval of 2 minutes. DIC/phase contrast and GFP fluorescence time lapse images were captured before and after stimulation with immunogenic dsDNA. For high frequency time-lapse imaging, we used a Fluoview FV1000 confocal microscope at a scan speed of 2 and sampling interval of 0.5 seconds. Frame sizes are indicated by scale bars. For intravital imaging, dorsal window chambers were installed in adult male reporter mice (8 to 24 weeks old). After window stabilization, 1 million MC38-H2B-mCherry colon cancer cells were injected into the tissue underlying the window and imaging was performed at serial time points.

### Quantitative Analysis of Epifluorescence Imaging

Maximum intensity projections of fluorescence images were used to define regions of interest (ROIs) for each single cell. For each ROI, calcium fluorescence was quantified across time, normalized to median of fluorescence trace, and expressed as ΔF/F = (F(t)-F(*t*_*median*))/F(*t*_*median*). Cell viability was quantified based on microscale changes in differential DIC images defined as DIC(t) – DIC(t-1). Pixels of each differential DIC image were summed, resulting in a scalar at each time point that reached a minimum that was interpreted as the timing of cell death. Cells were aligned by setting the timing of peak calcium fluorescence to zero.

### Inference of Cell Communication from Calcium Reporter Time Lapse Imaging

All analyses were performed using ImageJ/Fiji and MATLAB(Mathworks). Single cell regions of interest (ROI) were defined using ImageJ/Fiji and single cell fluorescence versus time was extracted for each ROI in all data except Figure 6. Background was corrected using a rolling ball algorithm and average fluorescence was quantified for each ROI across time. Each ROI fluorescence time-series was smoothed using a zero-phase digital low pass infinite filter which strengthens the passband signals with a cutoff of 0.45 Hz half power frequency. Peak-finding was performed and transformed into unit magnitude impulses located at the time of each fluorescence peak and termed the time of spike (TOS). This allowed construction of a discrete time series spike train *S*_*i*[t] for each cell *i*. Cross correlation plots were generated by rank ordering all calcium spikes of all cells within an individual movie. For each pair of calcium spike pair, we calculated the product of the difference in spike timing ΔTOS and the Euclidean distance raised to the 1.5 power ΔD^1.5^.

### Spatiotemporal Excess Synchrony pipeline details

Methods for temporal excess synchrony calculation are detailed in the primary manuscript text. Spatiotemporal synchrony was calculated as (ΔS/w) for each 50μm × 50μm sub-image, containing an average of 11 active ROIs, at moving temporal windows of size w. Regions of excess synchrony that were connected in either time or space were combined and interpreted as a single cell communication event. Synchrony “stringency” was defined a margin above S_th_ that would be interpreted as a genuine cell communication. It was expressed as a fraction of S_max_-S_th_ for each movie and therefore ranged between 0 and 1.

### Statistical analysis

All statistical analyses were conducted with GraphPad Prism software and MATLAB. Data are presented as mean ± SEM. Statistical significance was evaluated using the two-sided Mann-Whitney test or Kolmogorov-Smirnov test. P values less than 0.05 were considered to denote significance.

**Supplemental Figure 1.**
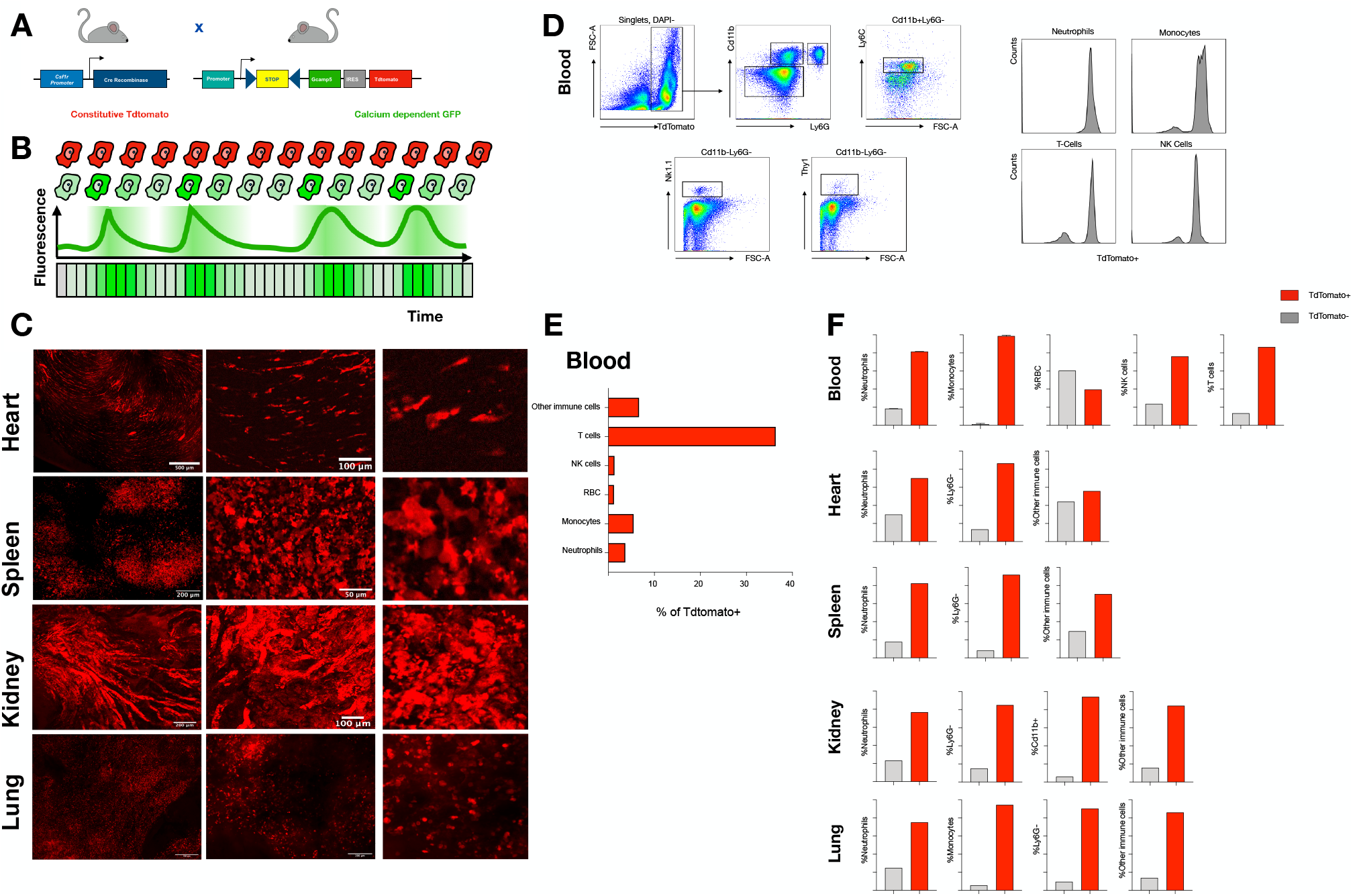
Design, construction, and characterization of a Csf1r-Gcamp5 calcium reporter for non-destructive quantification of innate immune cell dynamics. A) Illustration of mouse breeding strategy. Csf1r-cre mice were crossed with GCaMP5 inducible reporter mice to create an innate immune cell specific reporter. B) Cartoon illustrating that tdTomato is constitutively expressed as a reference and dynamic calcium-dependent GFP signals are quantified ratio-metrically C) Spatial distribution of tdTomato+ cells in solid organs (heart, spleen, kidney, lung) at low (left), medium (middle), and high (right) magnification (scale bars from left to right, 500**μ**m, 100**μ**m and 10**μ**m). D) Gating strategy for flow sorting of immune subsets. E) Percentage of total tdTomato^high^ cells in the peripheral blood from the Csf1r-GCaMP5 mouse. F) Percentage of each subset in each tissue compartment that is tdTomato+.

**Supplemental Figure 2.**
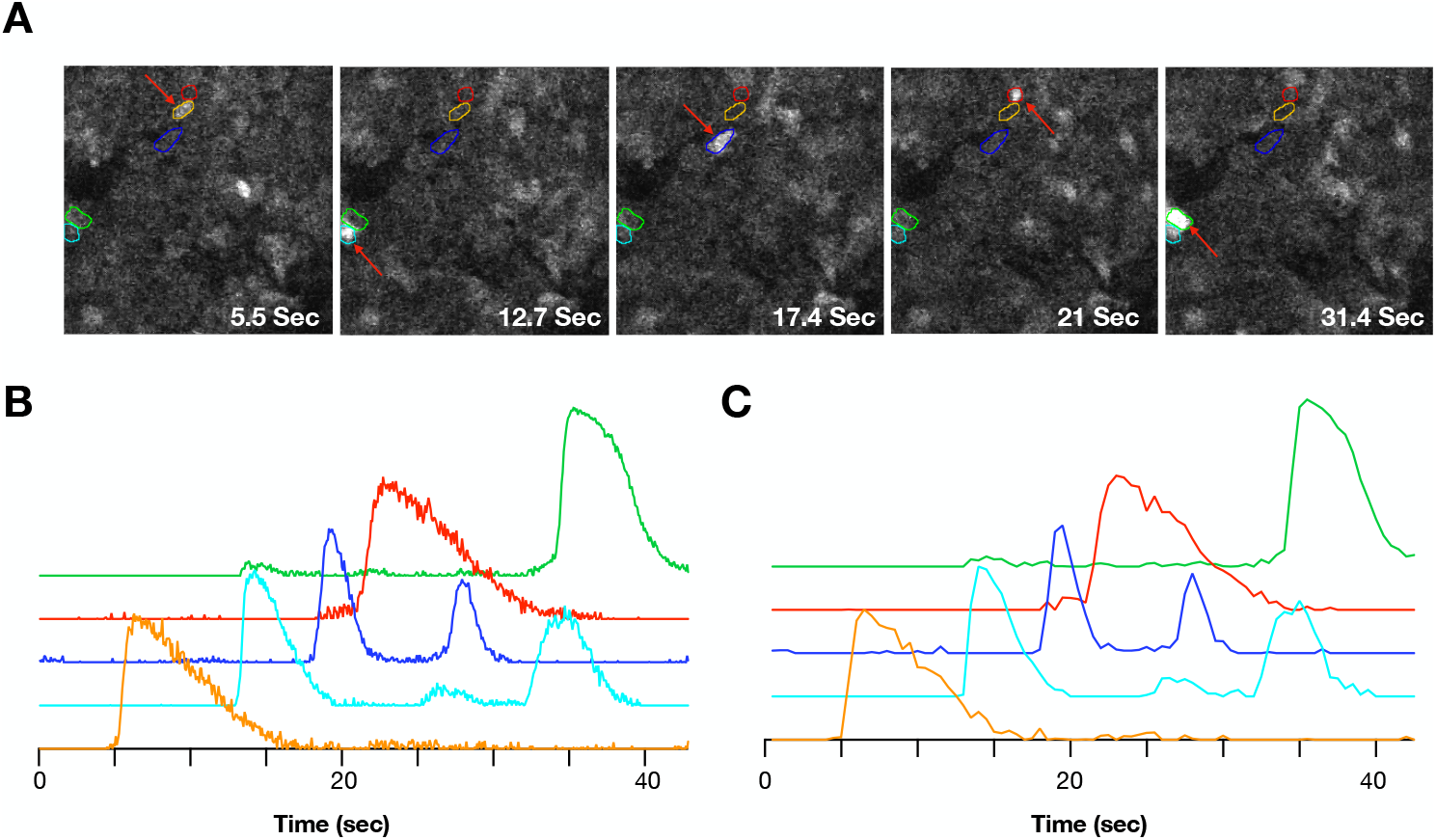
Csf1Cre-Gcamp5 sampling frequency determination. A) Calcium spikes were recorded from a population of cells at 15Hz. B) Quantification of calcium fluorescence versus time for multiple cells across time at 15Hz. C) Down-sampling was performed and every 7^th^ sample was plotted to show the similarity of calcium spike tracings. This led to selection of 2Hz as the sampling frequency used throughout the manuscript.

**Supplemental Figure 3.**
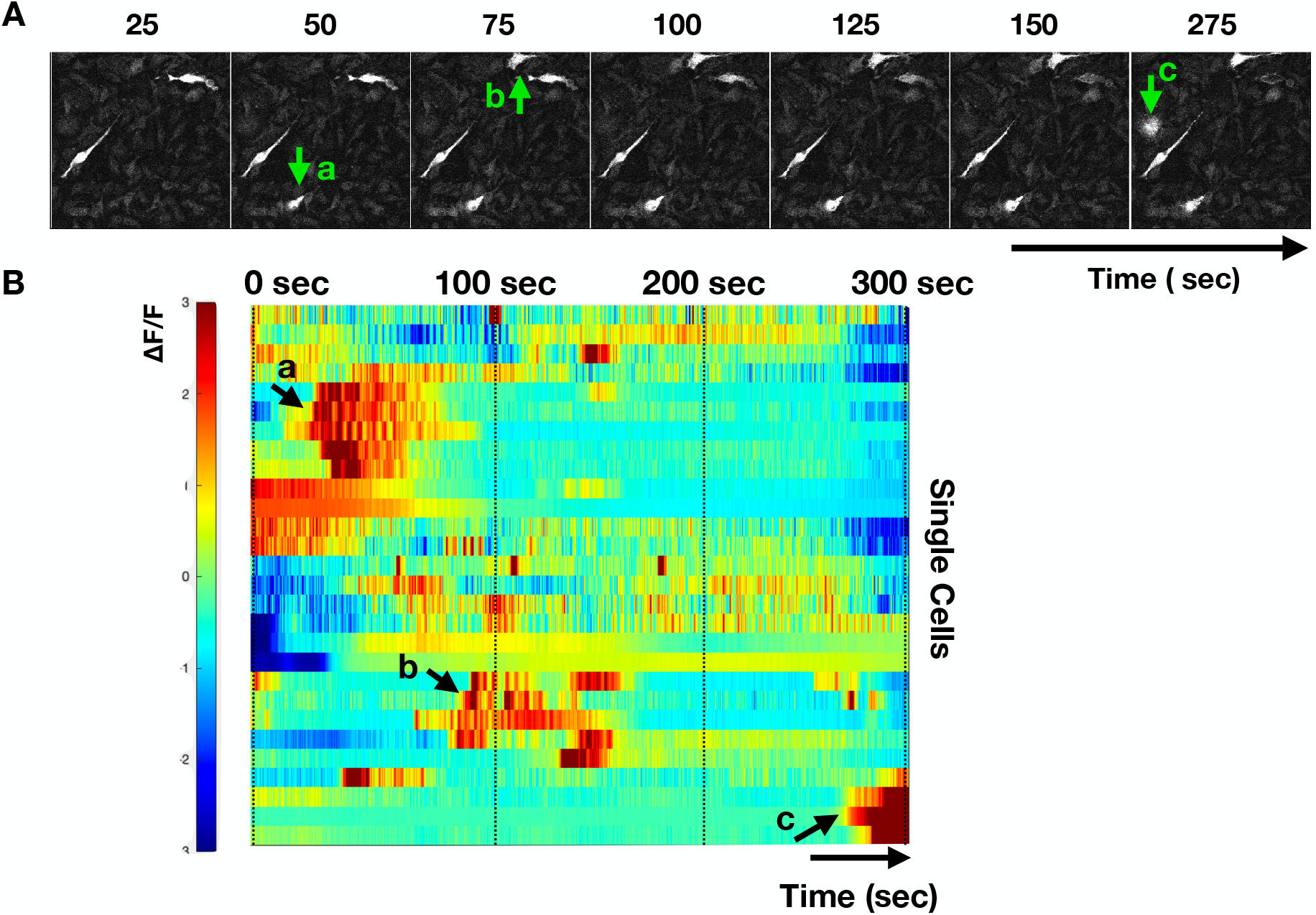
Example of Csf1Cre-Gcamp5 macrophage calcium reporter dynamics following immunogenic double-stranded DNA stimulation in vitro. A) Montage of time-lapse imaging. Newly calcium-overloaded macrophages indicated in a, b, and c precipitate non-fatal calcium fluctuations in neighboring macrophages. B) Heatmap illustration of hierarchically clustered macrophage dynamics.

**Supplemental Figure 4.**
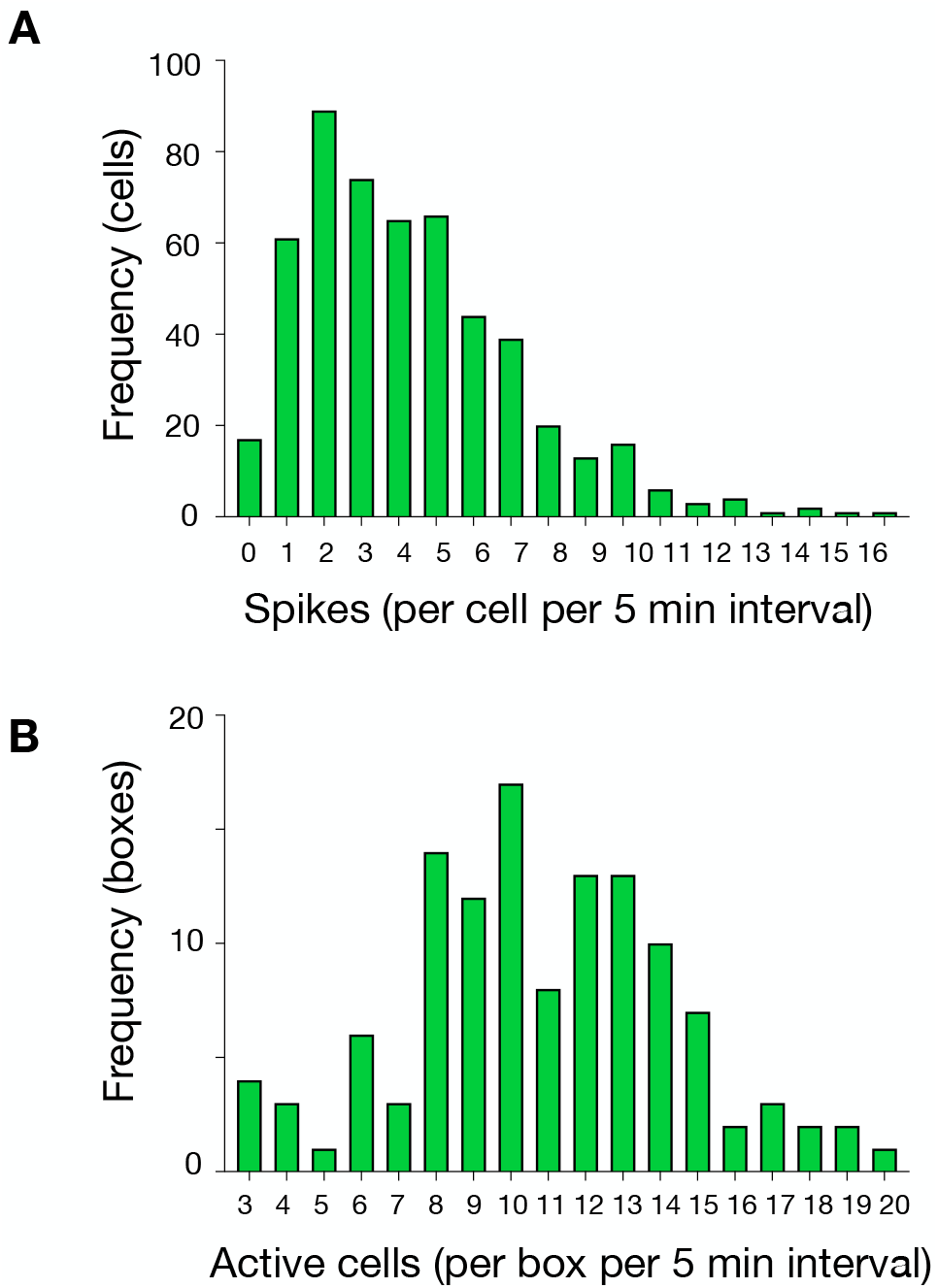
Histograms of in vivo Csf1Cre-Gcamp5 calcium reporter dynamics. A) Histogram illustrating distribution of number of spikes per cell per 5-minute interval. B) Histogram illustrating distribution of number of active cells per box per 5-minute interval.

#### SUPPLEMENTAL MOVIES

**Movies M1 and M2:** Time-lapse imaging of Csf1r-GCaMP5 bone marrow derived macrophages (BMDMs) measured at a 2 minute sampling interval on an Olympus Vivaview incubated epifluorescence microscope. (**M1**) Vehicle control stimulation. (**M2**) Stimulation with complexed immunogenic double-stranded DNA.

**Movie M3:**Time-lapse imaging of DNA-stimulated Csf1r-GCaMP5 BMDMs on an Olympus FV1000 confocal microscope oversampled at 15 Hz.

**Movie M4 and M5:** Time-lapse imaging of DNA-stimulated Csf1r-GCaMP5 BMDMs on an Olympus FV1000 confocal microscope, sampled at 2 Hz. (**M4**) Vehicle control stimulation. (**M5**) Transfection of complexed immunogenic double-stranded DNA.

**Movie M6-M9:** Intravital imaging (2 Hz, FV1000) of Csf1r-GCaMP5 reporter mice with a dorsal window chamber and an MC38-H2B-mCherry tumor. (**M6**) A discrete cell communication event occurs between 82-103 seconds in the lower left corner. (**M7**) A region of high frequency calcium spiking is shown. (**M8-M9**) In vivo dynamics of Csf1r-GCaMP5 reporter cells amidst MC38-H2B-mCherry tumor cells in a dorsal window chamber.

